# Correcting Methylation Calls in Clinically Relevant Low-Mappability Regions

**DOI:** 10.1101/2021.10.04.463127

**Authors:** Caiden M. Kumar, Devon P. Ryan, Bradley W. Langhorst

## Abstract

DNA methylation is an important component in vital biological functions such as embryonic development, carcinogenesis, and heritable regulation. Accurate methods to assess genomic methylation status are crucial to its effective use in many scenarios, especially in the detection and diagnosis of disease. Methylation aligners, such as Bismark and bwa-meth, frequently assign significantly higher MapQ values than can be supported by the uniqueness of the region reads are mapped to. These incorrectly high MapQs result in inappropriate methylation calling in repetitive regions. We observe reads that should map to separate locations (possibly having different methylation states) actually end up mapping to the same locus, causing apparent mixed methylation at such loci. Methylation calling can be improved by using Bismap mappability data to filter out insufficiently unique reads. However, simply filtering out Cs in insufficiently unique regions is not adequate as it is prone to over-filtering Cs in small mappability dips. These Cs can in fact often be called using reads anchored in a nearby mappable region. We have created a new feature for the MethylDackel methylation caller to perform read-based filtering. This new methylation calling method resolves some of the apparent mixed methylation to either 0% or 100% methylation and removes many unsupportable methylation calls. We examined methylation calls with and without read-based filtering in or near the 7830 genes containing ClinVar variants in a methylation sequencing data set from the NA12878 cell line. Use of this improved method corrected 41,143 mixed methylation Cs to 0% methylation, and 22,345 to 100% methylation throughout the genome.

## Introduction

As DNA methylation status can have a significant biological function [35], it is important that there be an accurate way of calling methylation on a genome. Although there are multiple varieties of DNA methylation, perhaps the most significant type in eukaryotes is methylation of cytosine to 5-methylcytosine [3]. Data on DNA cytosine methylation state can be gathered using a methylation sequencing technique (Figure 1), like bisulfite sequencing [5]. In bisulfite sequencing, unmethylated cytosines are deaminated to uracil by the addition of sodium bisulfite. 5-methylcytosines are not affected. Since uracil sequences as thymine and 5-methylcytosine sequences as cytosine, positions of unmethylated Cs in a reference sequence can be identified by C->T transitions[5].

**Figure 1:**
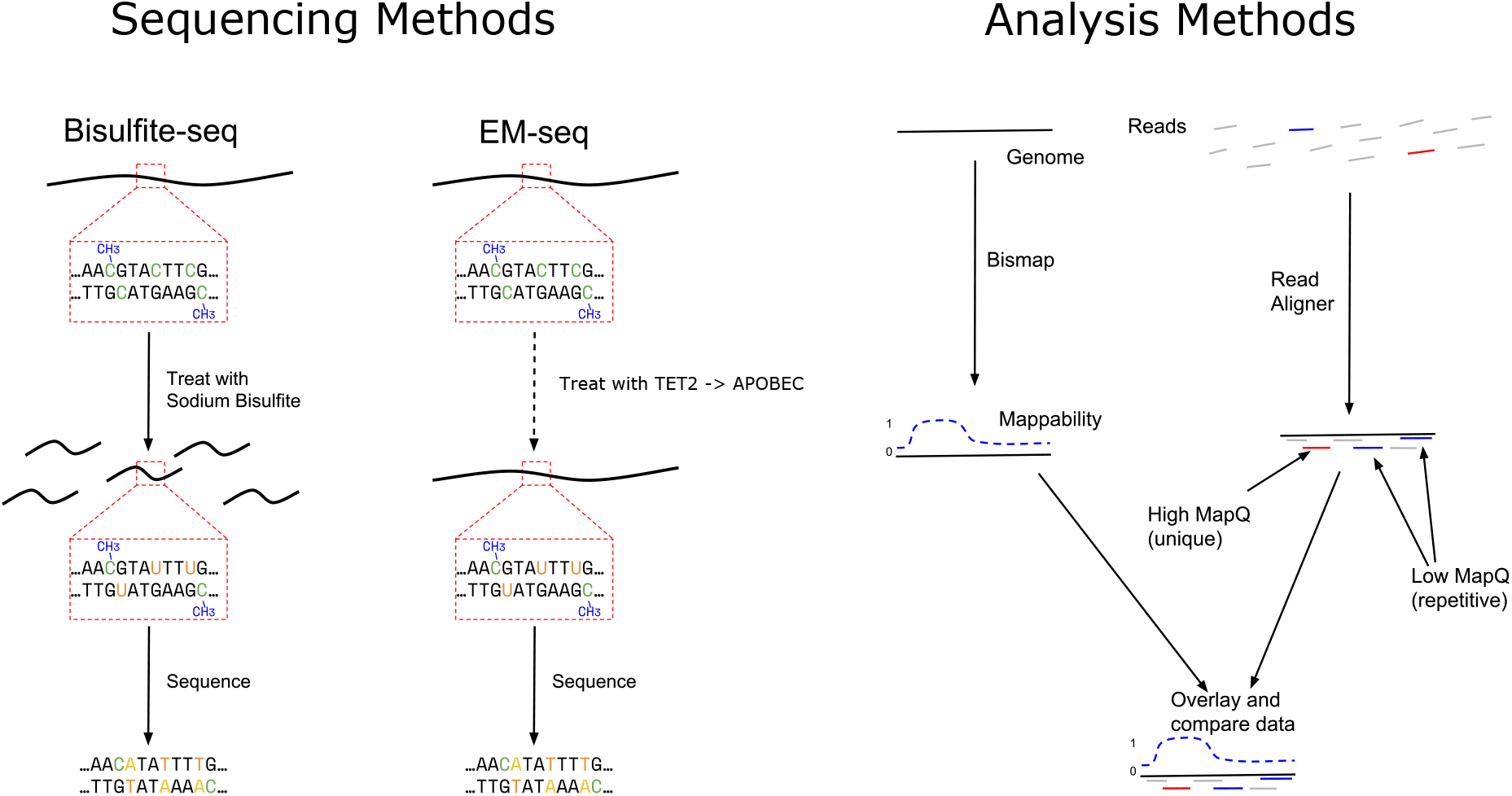
Overview of Methylation Sequencing and Data Analysis Methods.

It is also possible to use an enzymatic method, EM-seq, employing TET2 to oxidize 5-methylcytosine and an APOBEC enzyme to deaminate unmodified cytosines to uracil. While sodium bisulfite treatment produces double strand breaks and other DNA damage, this enzymatic method deaminates with more precision [38] and longer, better mappable fragments.

Whichever method is used to deaminate cytosines, methylation sequence data is typically aligned to a reference genome using a methylation-aware aligner [9], which is specifically designed to handle the C->T transitions in methylation sequencing data when aligning the reads to a reference. Once aligned, a methylation caller is needed to analyze alignments and report the methylation status of particular cytosines (Figure 1). The resulting data shows the methylation status of each callable cytosine in the genome and can therefore be used to find and study biologically significant DNA methylation sites.

There are two commonly used methylation callers, MethylDackel [34] and bismark_methylation_extractor [19]. The Bismark _aligner _creates XM, XR, and XG tags that are used by the bismark_methylation_extractor to call methylation. Bismark_methylation_extractor requires these larger bam files and is not compatible with bam files produced by other aligners. MethylDackel extract is not tightly coupled to a specific aligner or tag scheme, performing the filtering, quality assessment, and reference comparisons needed for methylation calling internally. MethylDackel efficiently loads a complete or targeted reference genome, examining alignments in chunks that are processed in parallel with all the context needed to differentiate methylation from C->T mutation. Flexible and optional thresholds are available for base-quality, positional trimming, adjacent base context (e.g. CpG or CHH), alignment flags, and MapQ.

A read must be unambiguously placed if it is to provide information about a specific region of a genome. Reads that equally match more than one area of the reference genome should not be used to assess methylation of any given C. To avoid calling Cs using reads derived from multiple genomic loci, methylation aligners (and read aligners in general) assign a MapQ value to each read alignment (Figure 1). According to the SAM specification [11], MapQ is defined as: “−10 log_10_ Pr{mapping position is wrong}, rounded to the nearest integer”. MapQ indicates how uniquely placed a read alignment is, that is, in how many other places could the read align to the reference. A low MapQ means that the read may align in many places throughout the genome (for instance, a read of centromeric satellite DNA would likely have a very low MapQ). A high MapQ indicates that the read likely aligns where it is placed and nowhere else in the genome.

A methylation caller can use accurate MapQ values to filter out reads with multiple placements in the genome, allowing the resulting methylation calls to accurately reflect their specific loci. Unfortunately, current aligners frequently produce implausible MapQ scores.

In this paper, we describe the problem that low-mappability creates for methylation calling, the extent of miscalled methylation in real datasets, and our implementation of an improved method that measurably reduces incorrect mixed-methylation calls improving differential methylation assessment, the use of methylation as a biomarker for disease progression, and understanding of development processes.

## Results

While evaluating the methylation aligner bwa-meth [28], we observed a significant number of reads with unexpectedly high MapQ values in repetitive regions (e.g. centromeres, Figure 2). Another commonly used aligner Bismark[20] produced even more calls in repetitive regions (data not shown). After observing high MapQ reads in the centromere and larger repetitive regions, we investigated to see if smaller regions might also be too repetitive to support the high aligner MapQ estimates observed. We identified repetitive regions using data from Bismap [16], a tool that counts the number of occurrences of every single K-mer of a particular length (in this case, k=100) in the genome to create a mappability score (ranging from 0 to 1 in increments of 0.01) for every base in the GRCh38 reference. Each individual K-mer is considered either mappable or not, and the overall mappability score is derived (indirectly) from how many different K-mers are mappable and how many are not. Bismap takes the effect of C->T conversion into account and therefore produces data which is applicable in the context of methylation sequencing [16]. Reads entirely contained within a region of low mappability should not have high MapQ values due to their repetitiveness, however we observed many high-MapQ reads in regions with very low or zero Bismap mappability (e.g. Figure 3).

**Figure 2:**
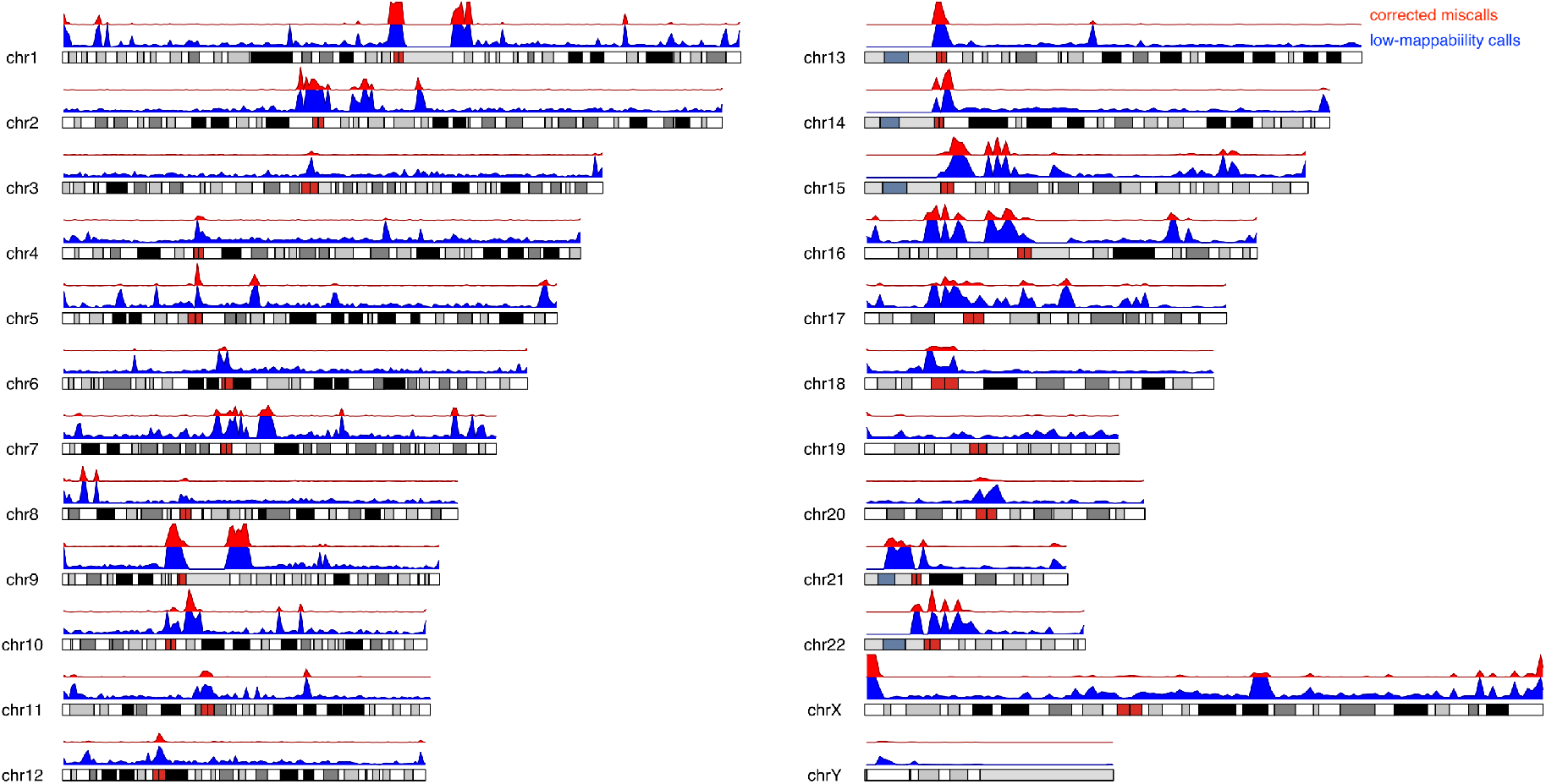
Cs in low mappability regions (blue), as well as those filtered out by per-read filtering that still passed MapQ filtering (red), are represented across all chromosomes. Per-read filtering removes unsupported calls while retaining millions of Cs that a nai”ve per-C filtering approach would needlessly eliminate. In this figure, the horizontal banded rectangles represent the chromosomes, the blue track above them shows all calls in low-mappability regions, and the red track corresponds to miscalls accurately removed by per-read filtering.

**Figure 3:**
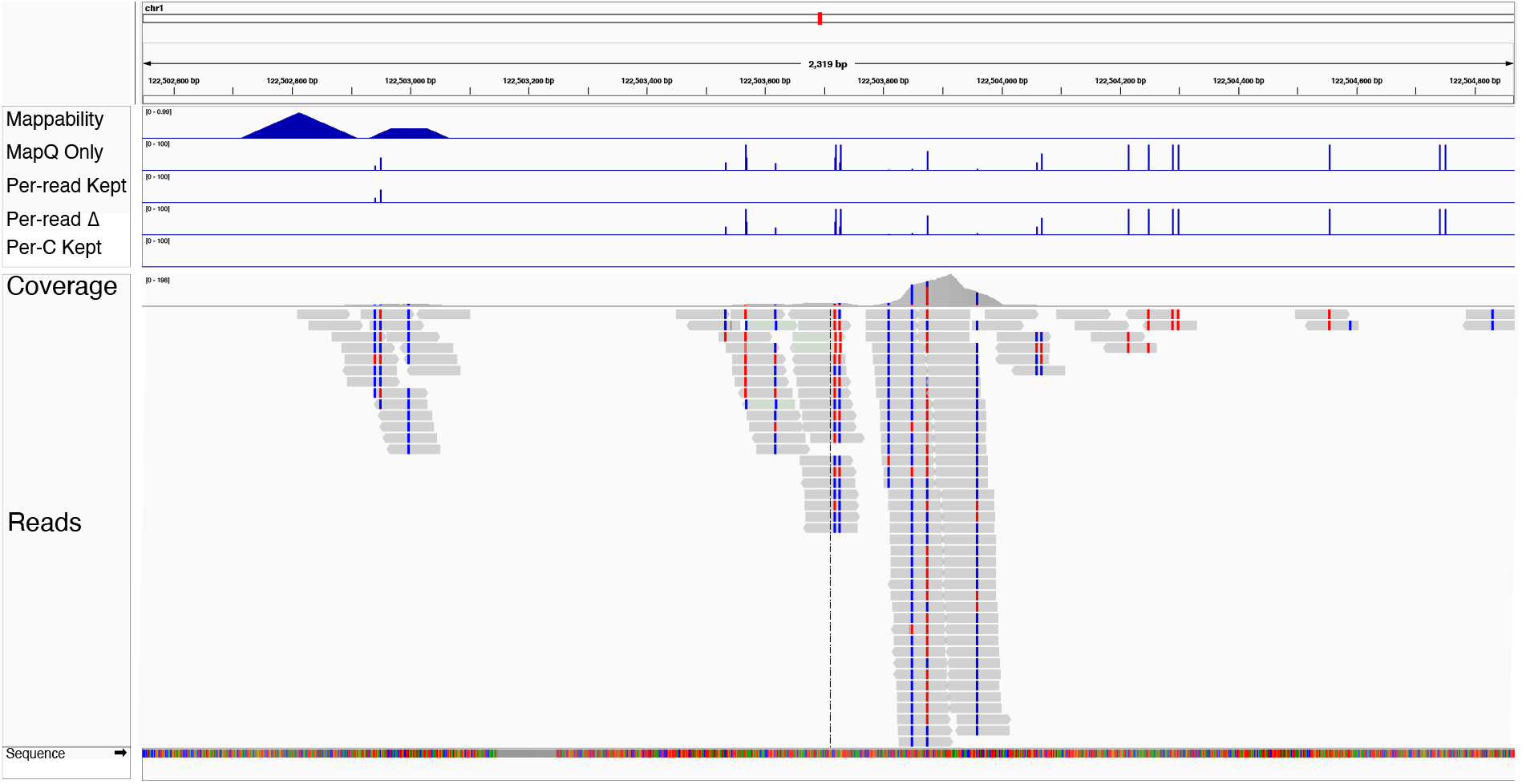
MapQ filtering does not remove all poorly mapped reads. This is an example of a coverage spike in a low-mappability region: clearly not sufficiently mappable, but still called with MapQ filtering. Mappability is only high in a small portion of this view (top track), but using MapQ alone alone enable calls of many Cs (MapQ Only track). the Per-read Kept track shows that only 2 sites have enough mappability to call and successfully eliminates the remaining miscalled Cs (Per-read Δ track). Per C filtering eliminates all Cs in this region including the 2 that should be callable (Per-C Kept track).

While low MapQ can indicate repetitiveness, even stringent MapQ thresholds cannot reliably select reads for safe methylation calling in regions containing repetitive DNA (Figure 4). We considered excluding methylation calls on Cs found in low-mappability regions (e.g. using bedtools), but rejected this approach because it is prone to both over- and under-filtering. Cs in short, unique regions would be kept (under-filtered) even if the surrounding DNA is repetitive (Figure 5). However, this situation was not commonly observed. Over-filtering of Cs in short repetitive regions is a larger problem though. In this scenario, a C located in a small dip in mappability would be eliminated (over-filtered) despite coverage from read pairs anchored in nearby unique regions (Figure 6). When calling using EM-seq reads aligned with bwa-meth to GRCh38, under-filtering is rare (between 9.4 * 10^−5^% and 1.2 * 10^−4^% of called Cs) but over-filtering is much more common (between 1.2% and 1.4% of called Cs) (Figure 4). Although ~1% of called Cs might seem like a small fraction, it accounts for ~14.0-16.2 million Cs, while ~10^−4^% of Cs corresponds to ~1079-1388 Cs. In cases where reads from multiple low-mappability regions are placed onto one region (creating a coverage spike), filtering by coverage could possibly also remove the problematic region, but in cases where reads are simply mixed among multiple low-mappability regions (not creating a coverage spike), coverage filtering would not be effective at removing the affected regions. In addition, if some of the reads used to call a C are uniquely mappable, but others are not, per-read filtering can remove only those reads that are not uniquely mappable, rather than being forced to discard the entire locus as is required for per-C filtering.

**Figure 4:**
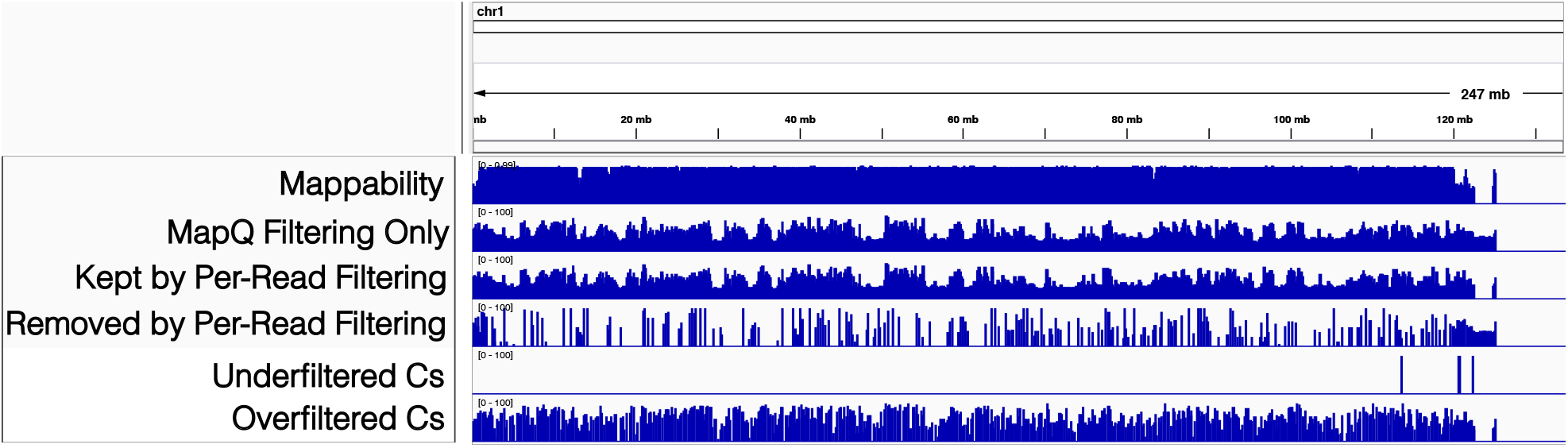
An overview of the impact of the various filtering scenarios in the q arm of GRCh38 chromosome 1. The six tracks show the effect of per-C and per-read mappability filtering on the methylation calls on this chromosome arm and are representative of rest of the genome. “Mappability” is the Bismap mappability for the chromosome. “MapQ filtering only” is the calls after commonly used MapQ filtering only (no per-C or per-read mappability filtering). “Kept by Per-read Filtering” shows the methylation calls kept after per-read mappability filtering. “Removed by Per-read Filtering” shows those calls removed entirely by per-read filtering for being insufficiently mappable. “Under-filtered Cs” are not sufficiently mappable according to per-read filtering, but are retained by per-C filtering. “Over-filtered Cs” are sufficiently mappable according to per-read filtering, but are removed by per-C filtering.

**Figure 5:**
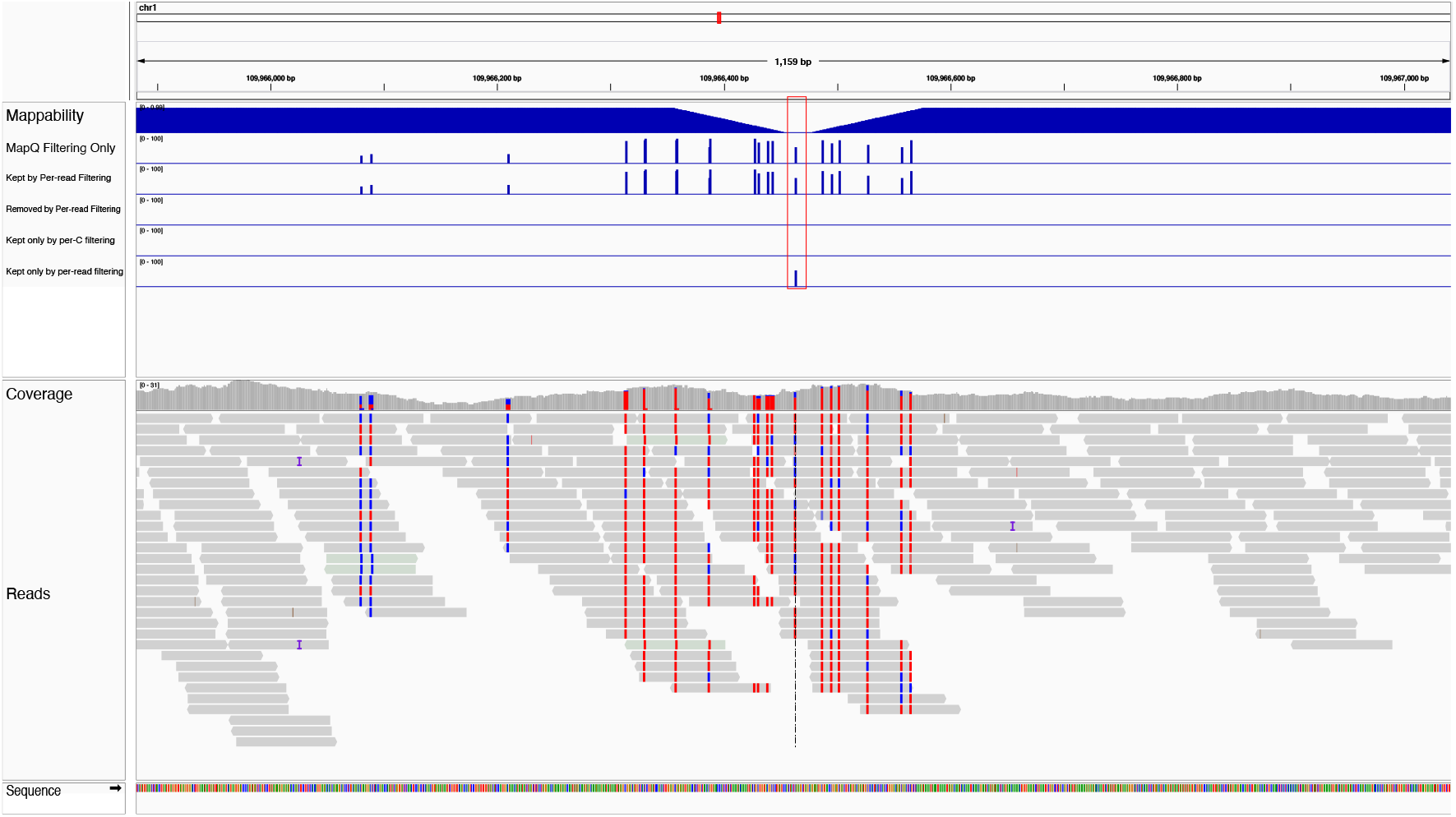
An example of a C incorrectly discarded by per-C filtering (red box). The C is in a small mappability dip, causing it to be discarded by bedtools filtering, while per-read filtering correctly retains the C, as it can be called using reads extending into the high-mappability regions on either side.

**Figure 6:**
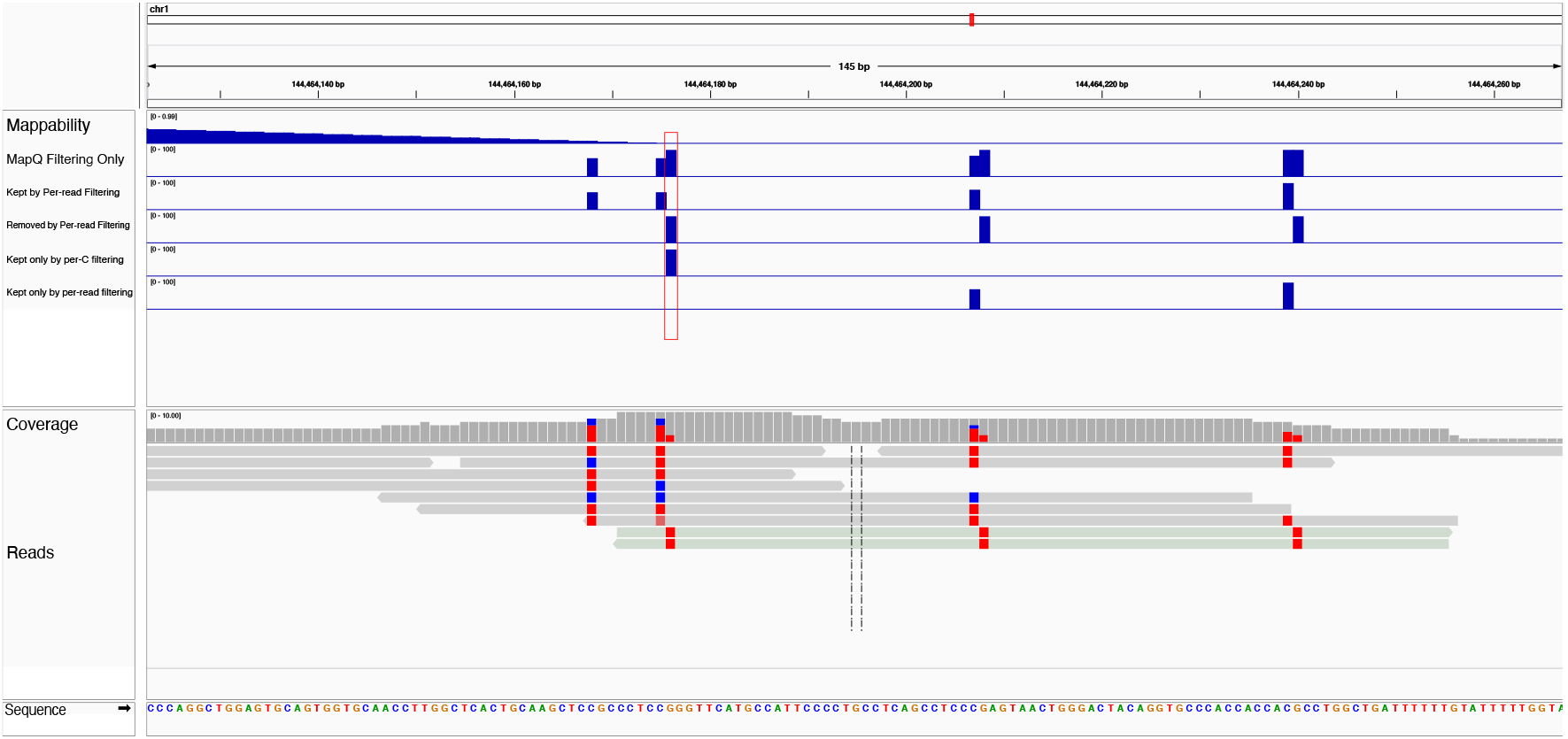
An example of a C incorrectly kept by per-C filtering (red box). As the C is itself just inside the edge of a mappable region, bedtools retains it, even though it is only supported by two insufficiently mappable reads. Our per-read filtering correctly removes this sort of C, as there is not enough data here to call it confidently.

To reliably filter out only problematic read pairs (those where both mates are placed in low mappability regions) we modified the MethylDackel methylation caller to accept a bigWig file of low-mappability regions to exclude from analysis. This per-read filtering approach precisely eliminates only those reads in repetitive regions and has a negligible effect on execution speed (see computational methods for details).

To focus on the incorrect alignments and methylation calls that have biological and medical significance, we examined methylation calls in and just upstream of GENCODE genes [12] that contain variants listed in the ClinVar [21] database of disease-associated variants. Using our alignment filtering approach we successfully avoided miscalling 264 thousand Cs in these important regions in a 200 ng EM-seq sample which have insufficient mappability to support methylation calling. Briefly, to identify the number of methylation calls correctly removed (miscalls), we took the list of unfiltered Cs for the regions we were interested in and removed from it all Cs we kept after per-read filtering. The result was a list of every C that was correctly filtered out (Figure 7).

**Figure 7:**
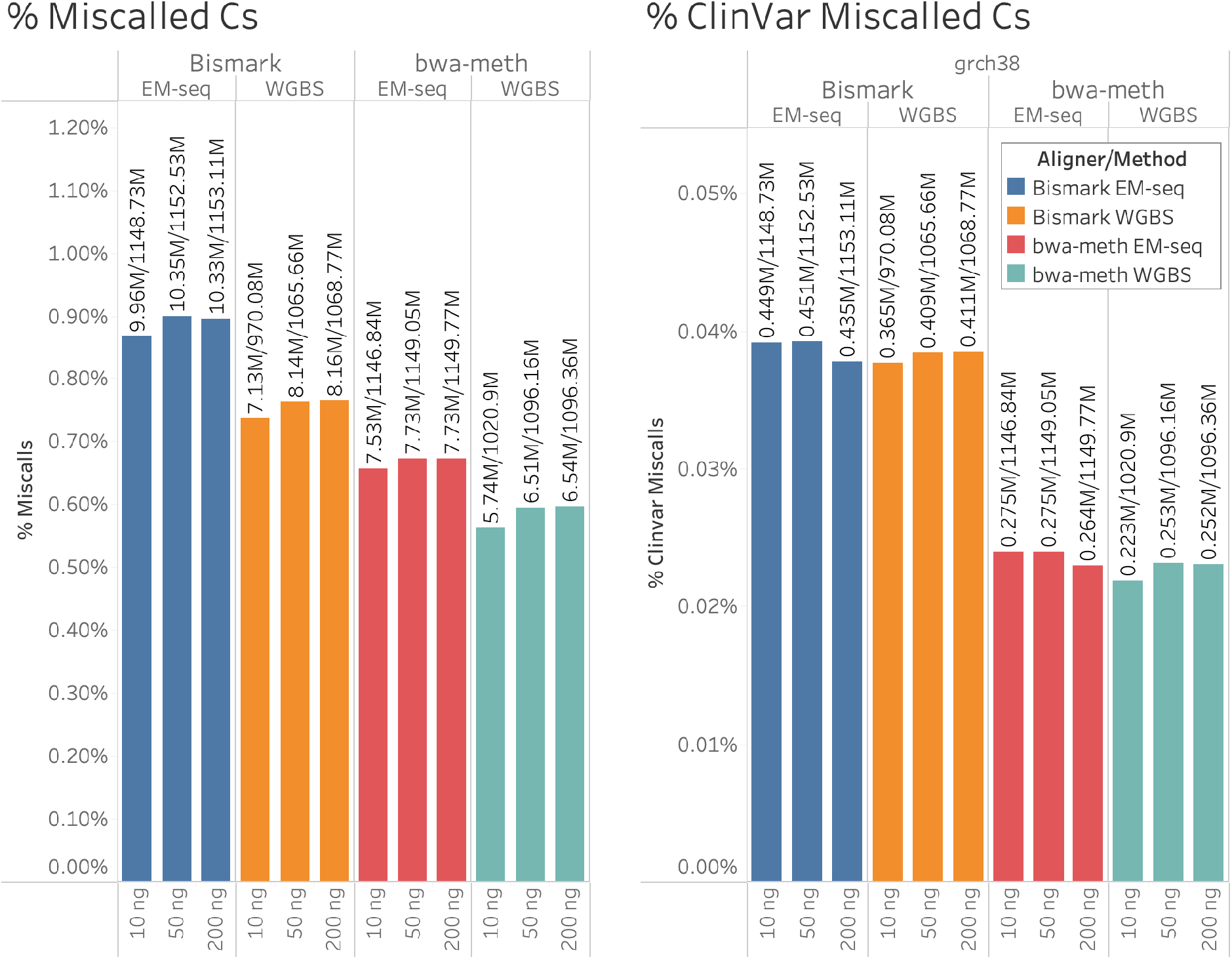
Left: Percents of miscalled Cs in the 10, 50, and 200 ng EM-seq and WGBS samples, aligned with bwa-meth and Bismark, where a “miscall” is a methylation call filtered out entirely by per-read filtering. The vertical axis is the percent of miscalled Cs in the sample. Right: Percents of miscalls within ClinVar genes in the same samples. The vertical axis is the percent of ClinVar miscalled Cs in the sample.

### Filtering Effect

We compared the effect of per-read mappability filtering on number of C’s called, as well as number of C’s called in low-mappability regions (as defined by per-C filtering), on EM-seq libraries of 3 sample masses aligned with the bwa-meth aligner. We saw a significant positive filtering effect on all 3 sample input amounts, with a mean of 7.67 million Cs filtered out by read-based filtering in each sample (see Figure 7). The miscalled Cs were scattered across the genome, as shown in Figure 2.

We also examined how many mixed methylation (0% < % Methyl < 100%) Cs were resolved to either 0% or 100% methylation, and found that approximately 60,000-90,000 methylation calls in the EM-seq data that were previously mixed were resolved by per-read filtering to either fully unmethylated or fully methylated (Figure 8), showing that unlike per-C filtering, per-read filtering can correct methylation calls without always having to entirely delete them.

**Figure 8:**
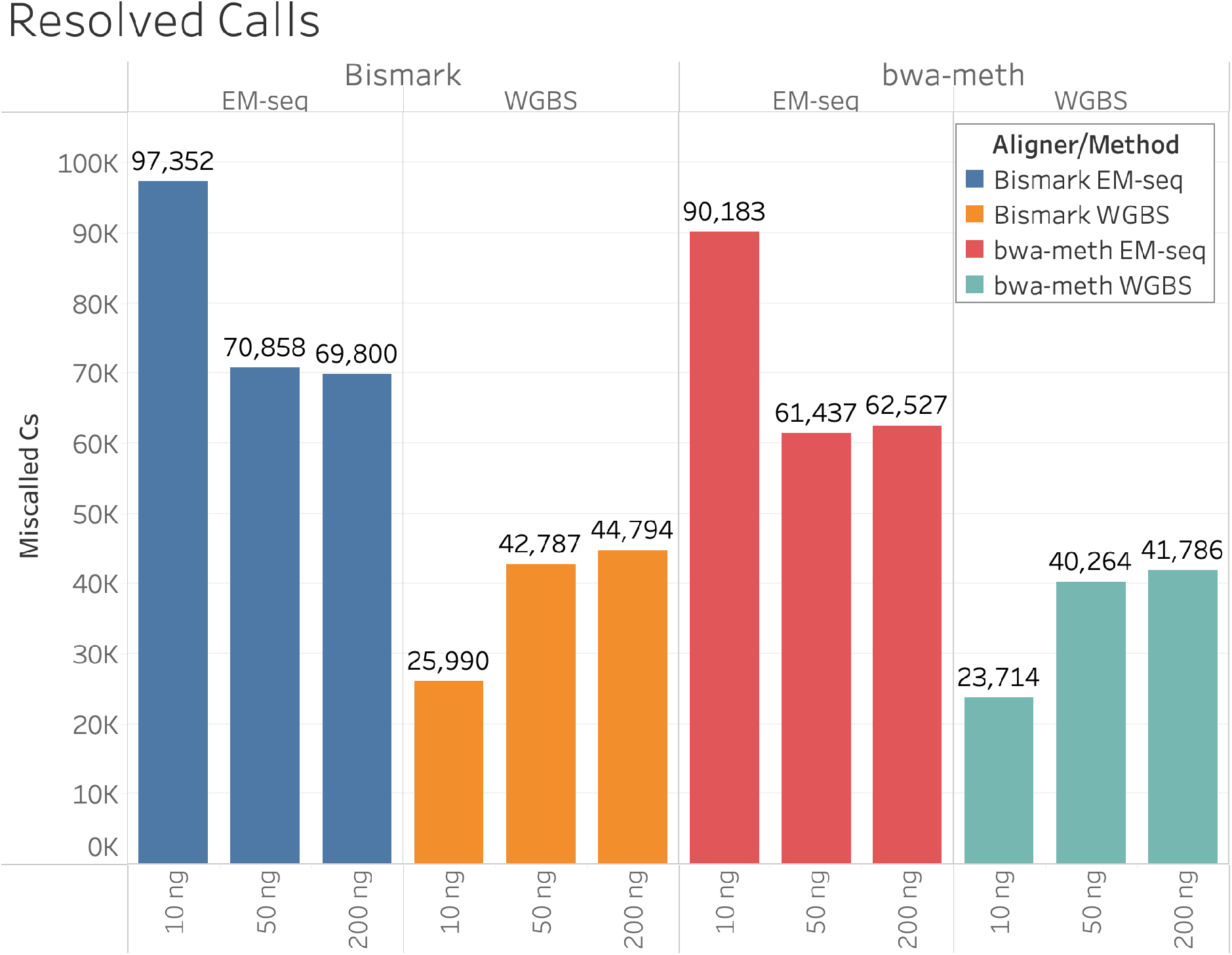
Counts of genome-wide resolved calls in the 10, 50, and 200 ng EM-seq and WGBS samples, aligned with bwa-meth and Bismark, where a “resolved call” is a methylation call that is mixed in the unfiltered data, but either 0% or 100% in the filtered data. The vertical axis is the number of resolved calls in the sample.

### Effect on Medically Relevant Genes

To estimate biological significance of these miscalls, we used genes in GENCODE containing variants in ClinVar (“ClinVar genes”). Focusing on genes that may contain miscalls, we filtered the list to only include those genes which had at least one region, either in the gene or in a 2 kb region upstream of the annotated start site (to capture regulatory regions), where the mappability is too low to allow reliable methylation calling. Examination of this data in IGV[33] and numerically revealed that there were miscalls in or upstream of thousands of unique genes in the EM-seq/bwa-meth data, with a median number of 64 miscalls per gene, indicating that indeed miscalls may affect biologically significant methylation. Many of these miscalls can be resolved to 0 or 100% methylation by the per-read filtering method described in this paper (Figures 8, 9).

**Figure 9:**
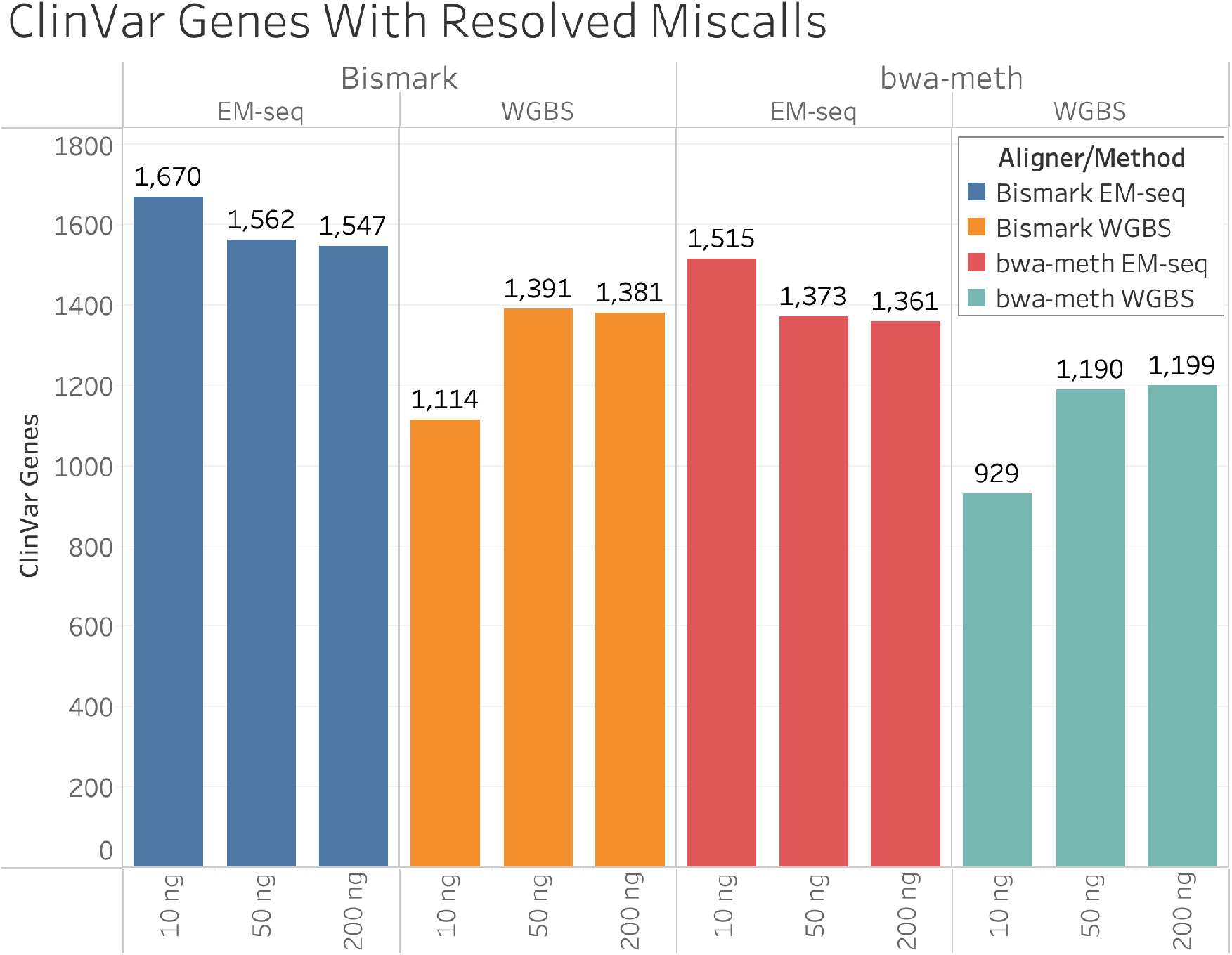
Counts of ClinVar genes containing resolved (from intermediate to 0 or 100% methylation) miscalls in the 10, 50, and 200 ng EM-seq and WGBS samples, aligned with bwa-meth and Bismark. The vertical axis is the number genes containing miscalls that were resolved by mappability-informed methylation calling.

It is evident that the sites of most methylation miscalls fall into two broad types. First, are repetitive regions that simply have a mappability of 0 for a long stretch of sequence. All Cs in such regions are classified as miscalls due to the lack of mappable regions to anchor reads with. Second, are shorter regions at the border between high mappability and low mappability regions. These regions are where per-C filtering produces over- and under-filtered Cs. Over-filtered Cs occur when a C is in the low mappability region, but there is a mappable read anchored in the high mappability region that allows it to be called. Under-filtered C’s occur when a C is just inside a mappable region, but there is no sufficiently mappable read to call it. Examples of both from a 200 ng EM-seq sample aligned to GRCh38 using bwa-meth are provided in Figures 10 and 11.

**Figure 10:**
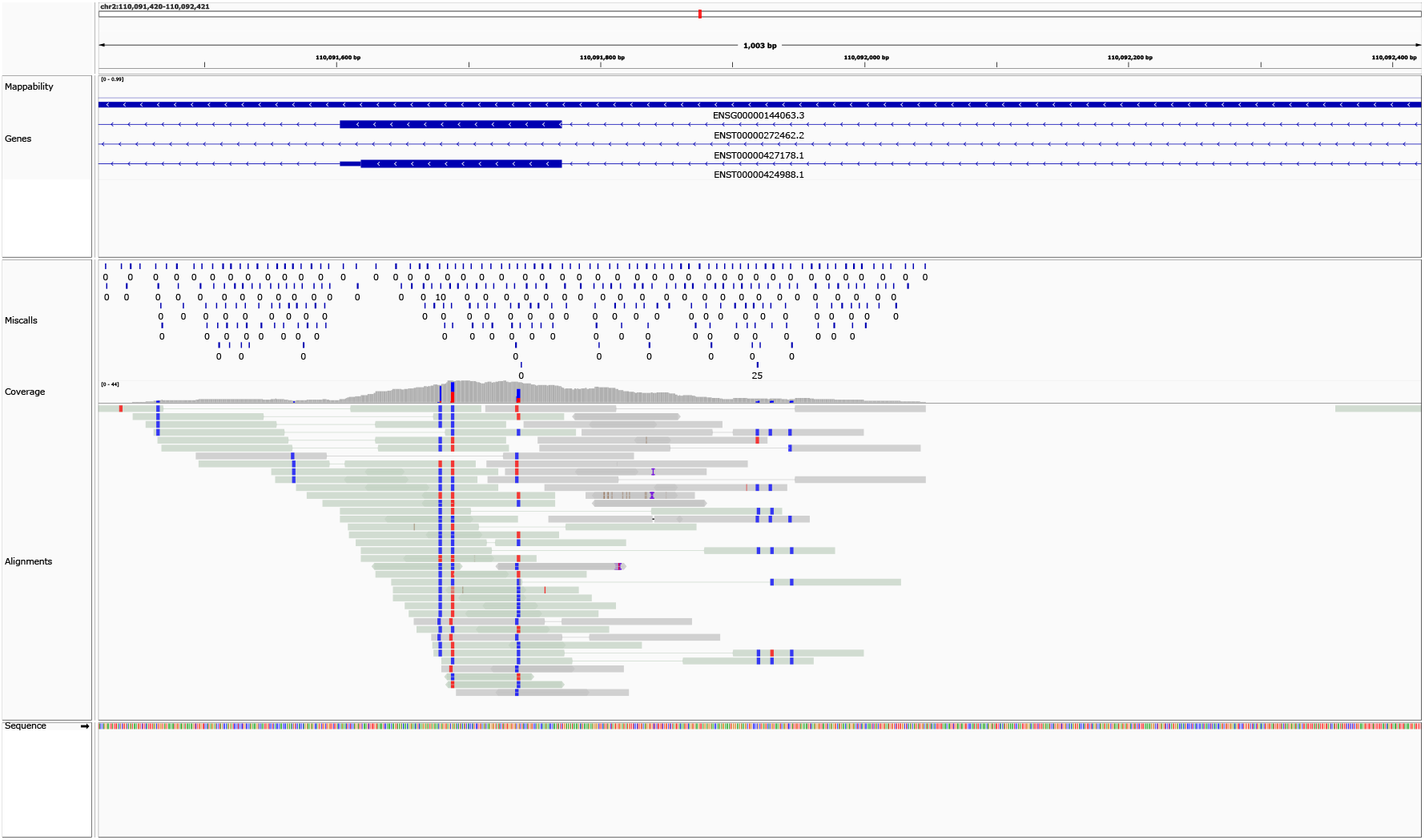
A view of a cluster of miscalls in a large low-mappability region containing the MALL gene (ENSG00000144063.3) [30], variants of which are associated with human disease[8], [15]. The reads are shown paired, with a line connecting reads 1 and 2 in a pair, with CpG methylation sites highlighted. Miscalls are shown in the track above the coverage graph. Note that every CpG call (as well as calls in other contexts) are miscalled in this region, as there are no high-mappability regions to anchor reads with. Two of these Cs show mixed evidence of methylation state, which may be due to DNA originating from multiple loci all mapping in this location with falsely high MapQ.

**Figure 11:**
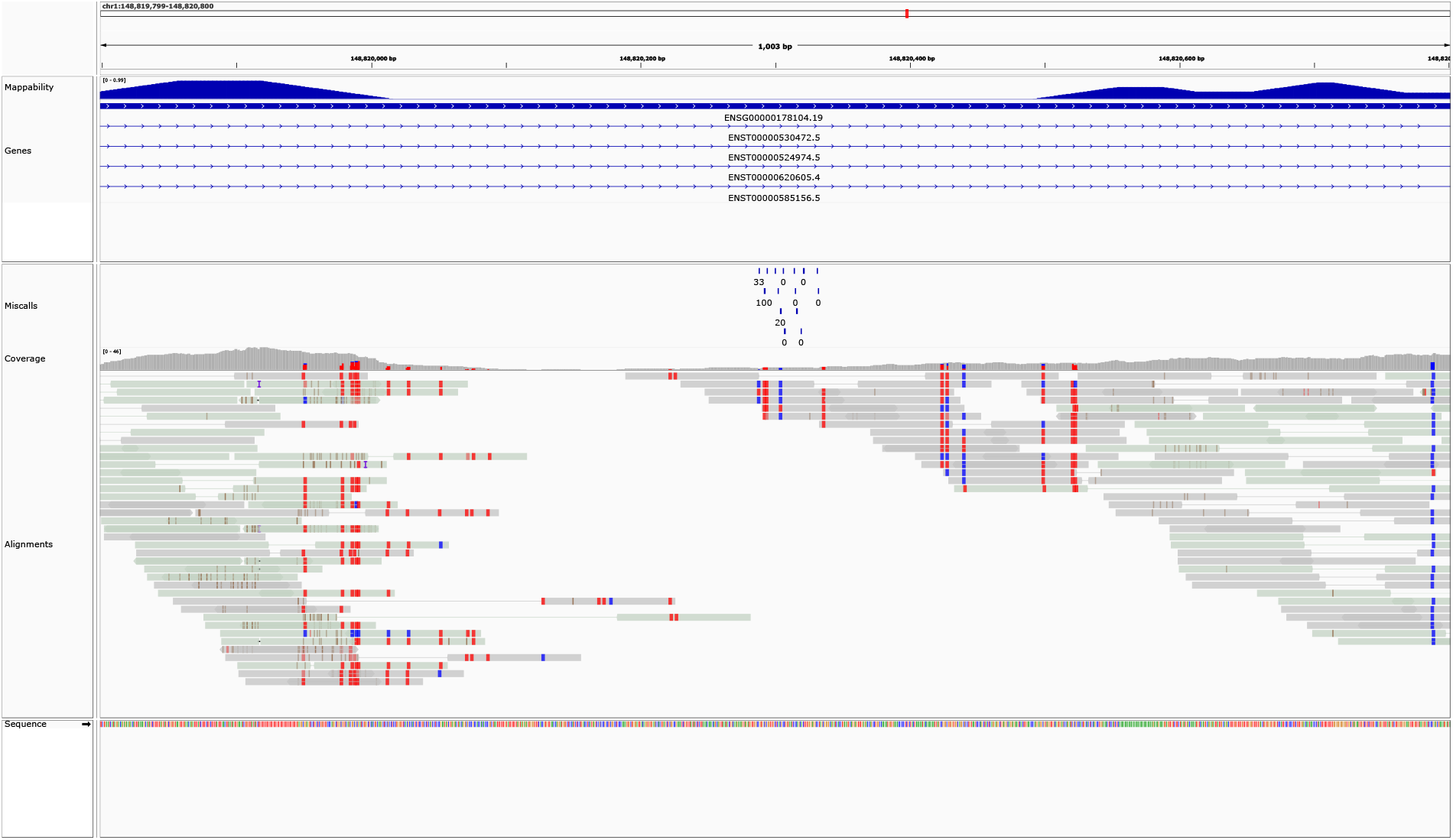
A view of a cluster of miscalls near the boundary between high and low mappability regions in the gene PDE4DIP (ENSG00000178104.19), regulation of which is reported to relate to dementia[29], schizophrenia[4], pineoblastoma[36] and other human disease[27, 2]. The read alignments are shown paired, with a line connecting reads 1 and 2, and highlighted for CpG methylation. Note that some methylation calls in the low-mappability region are not miscalls, as they can be called using read pairs extending into a high-mappability region.

### EM-seq vs. WGBS

Due to the reduced DNA damage and longer possible read lengths that are found in in EM-seq libraries, we compared with WGBS sequence libraries produced using the same 3 sample masses, to see if there is a difference between the two methods with regard to the number of miscalls. WGBS libraries have fewer overall Cs called[38] and therefor fewer miscalled Cs at all input masses than EM-seq libraries. In ClinVar gene regions, where both methods provide more similar overall coverage, the percent of miscalls is also similar between EM-seq and to WGBS libraries (Figure 7). We expected that longer reads could allow for more low-mappability Cs to be called using reads and mates extending into higher-mappability regions. Importantly, we observed that EM-seq libraries are able to resolve many more miscalled Cs throughout the genome (Figure 8) and in ClinVar regions than WGBS libraries 9).

### Bwa-meth vs. Bismark

For comparison, we examined the Bismark methylation aligner [19] in the same manner as the bwa-meth analysis. We observed that compared to bwa-meth, Bismark produced more inappropriately-high MapQ reads in repetitive regions (Figure 2). In addition, Bismark and bwa-meth use different systems to define values for MapQ. Bismark (as a result of using Bowtie2 for alignment) reports MapQ based on number of reference mismatches, producing values between 0 and 42 [31, 14]. Bwa-meth (using BWA-MEM for alignments) follows the SAM specification in estimating probabilities in the MapQ field. Overall, we observed a higher percent of miscalls with Bismark than we did with bwa-meth, as shown in Figure 7. This difference was noticeable in both overall miscalls and miscalls in ClinVar genes.

### GRCh38 vs. T2T

We also examined the effect of using the T2T reference [26] in place of GRCh38, since the T2T reference is much more complete than GRCh38, including repetitive regions such as centromeres, telomeres, and rDNA, which could play a role in where methylation is called and what areas of the genome have unique enough sequence to support calling. We found that, as we expected, there were far fewer miscalls in the T2T data as compared to the GRCh38 data, with less than half the number of miscalls in T2T as in GRCh38. Considering both strands, total calls were roughly similar between the two reference sequences with more Cs available in the T2T reference. (Figure 12, left). Use of per-read filtering has a smaller effect with the more complete reference, but thousands of Cs are still affected (Figure 12, right). This indicates that using the T2T reference can allow for better calling of methylation with fewer miscalls, however without a high quality annotation available, this may not yet be a practical option.

**Figure 12:**
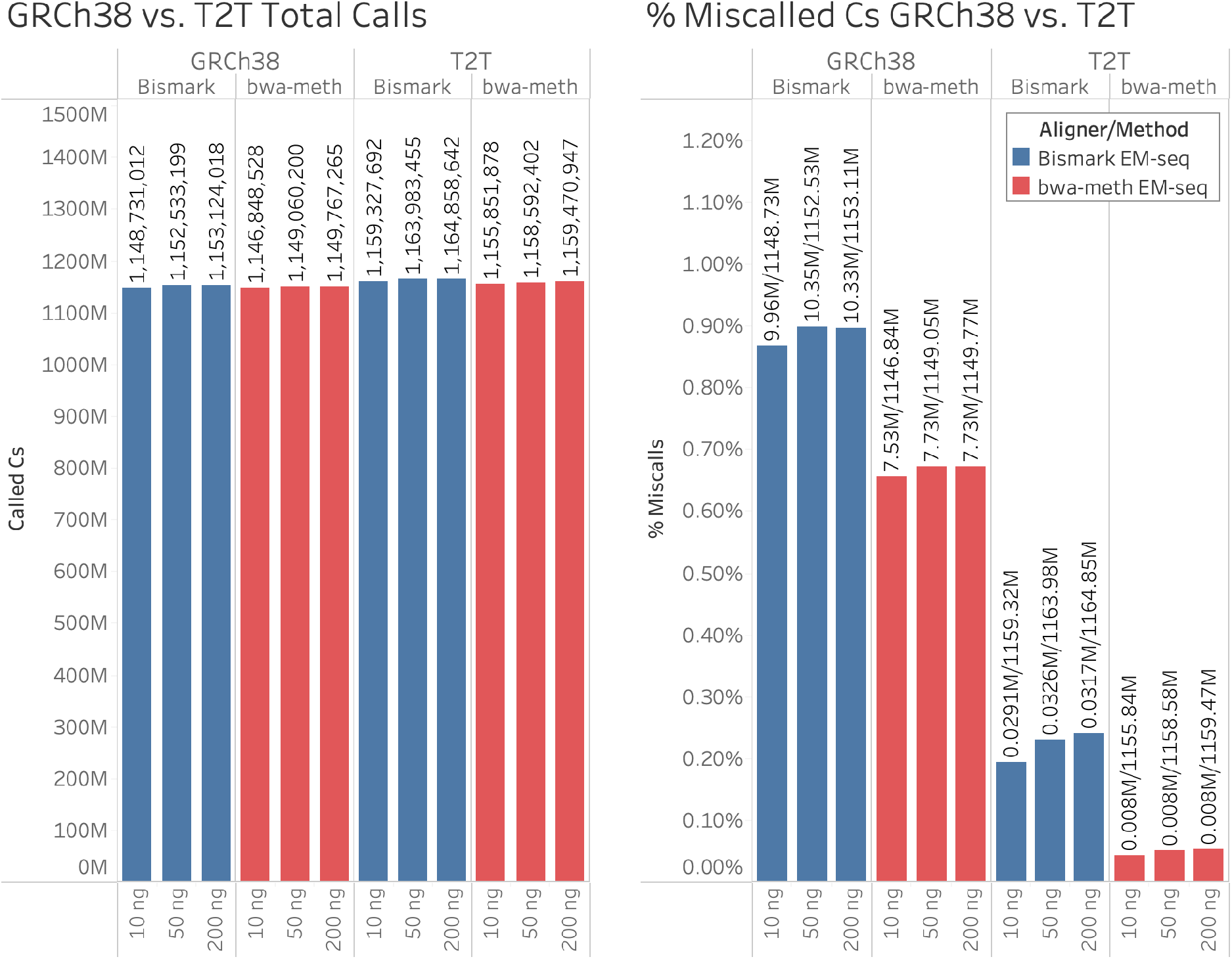
Left: Total call counts before filtering in the 10, 50, and 200 ng EM-seq samples, aligned with bwa-meth to GRCh38 and T2T. The vertical axis is the number of calls in the sample. Right: Percents of miscalled Cs in the same samples, where a “miscall” is a methylation call filtered out entirely by per-read filtering. The vertical axis is the percent of miscalled Cs in the sample. In both figures, the colors define which sequencing method and aligner were used, as detailed in the legend.

## Materials and Methods

### DNA Methylation Sequencing

#### Materials

Previously published[38] libraries from 10, 50, and 200 ng of genomic DNA from the NA12878 cell line (Coriel) were analyzed. This input DNA was supplemented with a small amount of lambda phage DNA (NEB #N3011), and a pUC19 plasmid (NEB #N3041) treated with M.SssI CpG Methyltransferase (NEB #M0226).

#### Whole Genome Bisulfite Libraries

The whole genome bisulfite (WGBS) libraries were prepared using the Ultra II DNA library prep kit (NEB E7645) before being Bisulfite converted using the Zymo EZ DNA Methylation-Gold bisulfite conversion kit according to the protocol described in Vaisvila et al. [38].

#### EM-seq Libraries

The EM-seq libraries were prepared using the NEBNext Ultra II DNA library prep kit (NEB E7645) (using the NEBNext EM-seq adapter), then EM-seq converted using TET2 and ABOBEC3A, as described in Vaisvila et al. [38].

#### Sequencing

Methylation libraries were sequenced with more diverse libraries (~10%) on 2 flow cells of an Illumina NovaSeq 6000 [13] using the S4 chemistry. We analyzed about 1.3 billion 99 bp paired-end reads for the for each library.

**Table 1:** Sequencing read counts after downsampling and adapter/quality trimming broken down by sequencing method and sample mass. Read counts are reported as counts of individual reads, not of pairs.

| Sequencing Method | Sample Mass | Read Count |
| --- | --- | --- |
| WGBS | 10 ng | 1,268,442,060 |
| WGBS | 50 ng | 1,269,601,144 |
| WGBS | 200 ng | 1,269,737,220 |
| EM-seq | 10 ng | 1,298,351,192 |
| EM-seq | 50 ng | 1,296,831,526 |
| EM-seq | 200 ng | 1,296,793,470 |

#### Computational Methods

The following data and tools were used for the analysis:

The GRCh38.p11 analysis set (hereafter referred to as “GRCh38”) supplemented with phage T4, phage lambda, phage Xp12, and pUC19 contigs was used throughout [6]. A VCF file of disease-associated variant sites was downloaded from ClinVar (see supplemental data for file date) and a GFF file of the GENCODE v31 gene annotations were used. The mappability data was the 100bp multi-read bigWig file downloaded from Bismap (see supplemental materials for link).

The Nextflow scripts used to analyze this data can be found at https://github.com/nebiolabs/low_bismap_methyl_calls. A detailed diagram of the analysis pipeline is found in Supplemental figures S.2 and S.3. Tools used include bedtools [32], samtools [22], GNU awk, MethylDackel, bigWigToBedGraph [17], BEDOPS [25], GNU sort, GNU head, bwa-meth, and Bismark. GNU sort and head are required since some of the functionality needed (specifically, the ability to parallelize sorting and the ability to use a negative value with the -n option for head to count lines from the end of the file) is not present in BSD sort and head.

## Discussion

Reads placed with falsely high confidence elicit cascading detrimental effects on methylation calling, differential methylation assessment, and assessment of phenotypes associated with methylation status. To remove the spurious, unsupported methylation calls that result from such reads, we suggest that reads with both mates in low mappability regions (as determined by Bismap) should be excluded from methylation calling. Additionally, because of the more accurate MapQ values, decreased run time, and more flexibility to separate methylation calling from alignment, we also recommend the use of bwa-meth for alignment and MethylDackel with MapQ > 10 and the new mappability feature enabled for methylation calling.

## Supplemental Materials

### Detailed Computational Methods

**Figure S.1:**
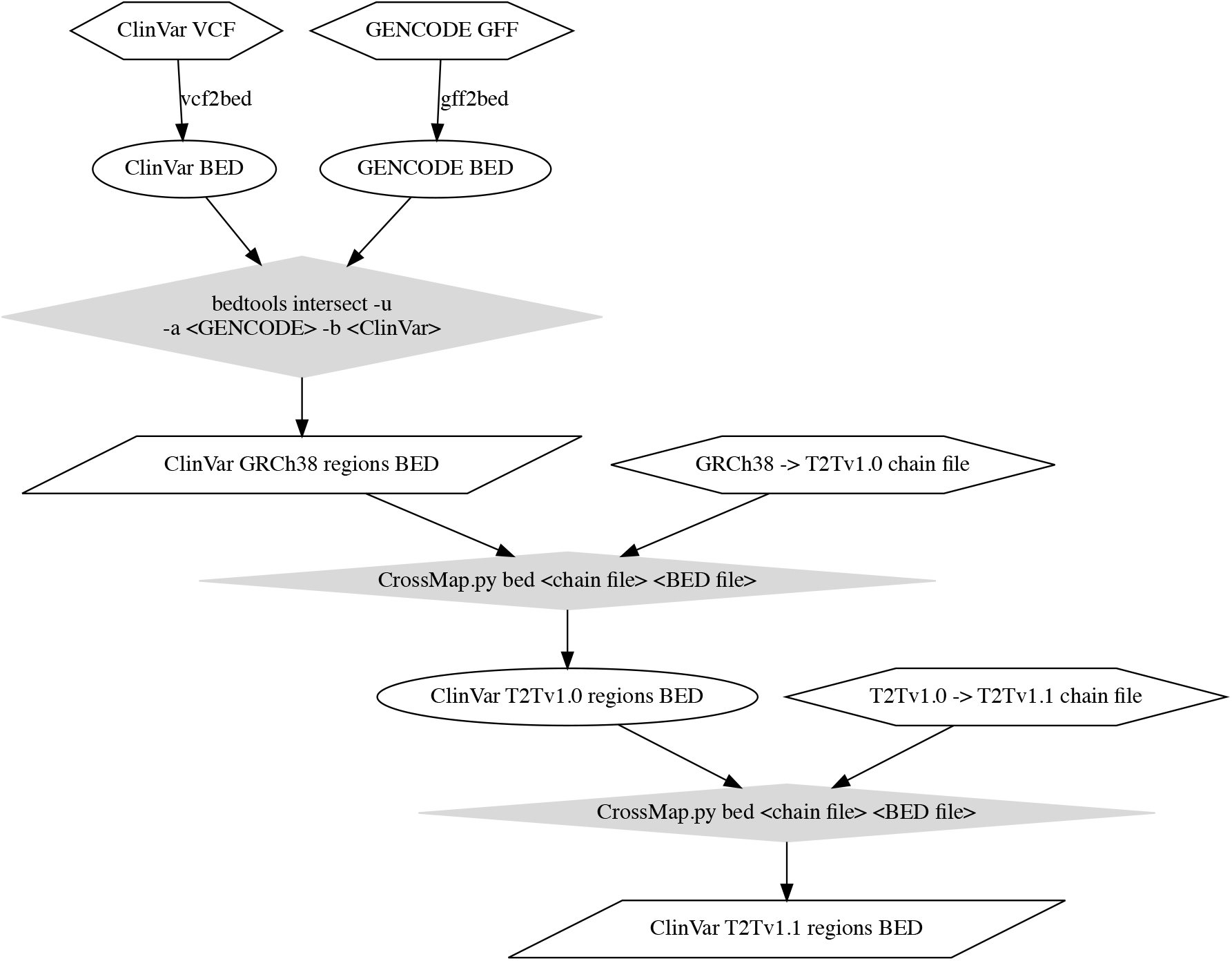
Detailed description of the procedure used to annotate the T2Tv1.1 reference using the GRCh38 ClinVar and GENCODE data. Hexagons are input files. Ovals are intermediate files. Gray rhombuses are processing steps with multiple arguments. Parallelograms are output files. Labels on edges are processing steps with one input and one output.

The sequencing reads were aligned using the bwa-meth aligner with default options. Methylation calling was performed on the resulting BAMs using MethylDackel v0.6.0. MethylDackel uses a default value of 10 as a minimum MapQ which should include mostly single locus reads, however inaccurate MapQs can lead to reads being incorrectly included in methylation calling. In order to eliminate reads in low-mappability regions, a patch was created for MethylDackel which allows it to take as an input a bigWig file which is then used to filter out read pairs (this patch currently only supports paired-end reads) where neither mate intersects a high-mappability region. The patch allows for user configuration of the low mappability threshold and the number of bases which must be equal to or above that threshold in order for the read pair to be kept (the defaults are a low mappability threshold of 0.01 and to require 15 bases that are greater than or equal to that threshold in a single read). The filtering algorithm has been optimized both by loading the mappability data into memory before calling, and through the use of a custom run-length compressed binary file format that we here term BBM (Binary BisMap). This format can store the mappability data for the hg38/GRCh38 human genome in 143 MB, compared to 1.11 GB when stored as a bigWig, giving a compression ratio for this dataset of 7.78:1. MethylDackel’s trimming options were set to trim off the first 1 bp of read 1 and the last 2 bp of read 2.

**Figure S.2:**
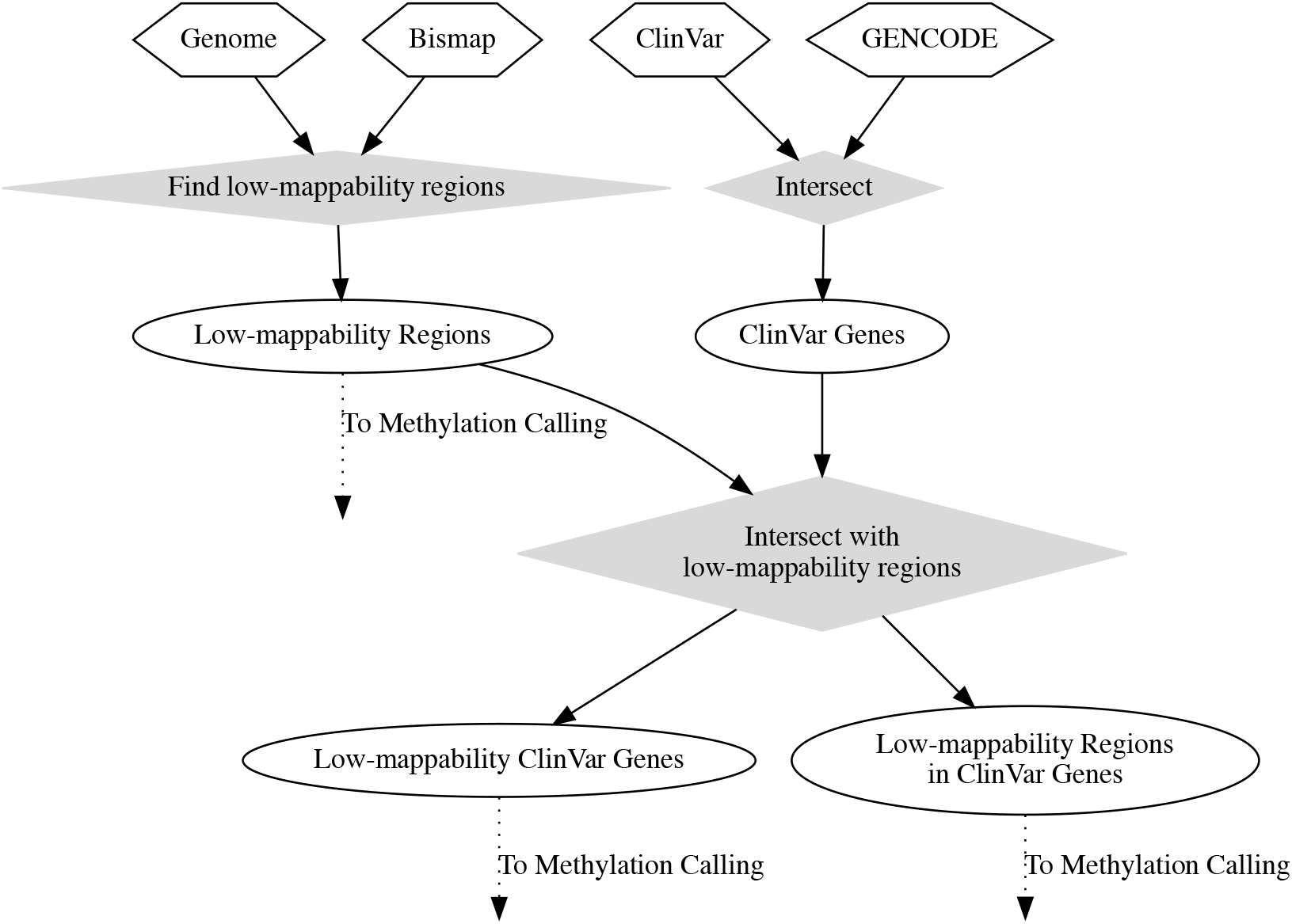
Detailed description of preprocessing steps used to identify biologically relevant low-mappability regions. Ovals are intermediate files. Gray rhombuses are processing steps with multiple arguments. Dotted lines indicate inputs that go to the methylation calling steps in Figure S.3.

**Figure S.3:**
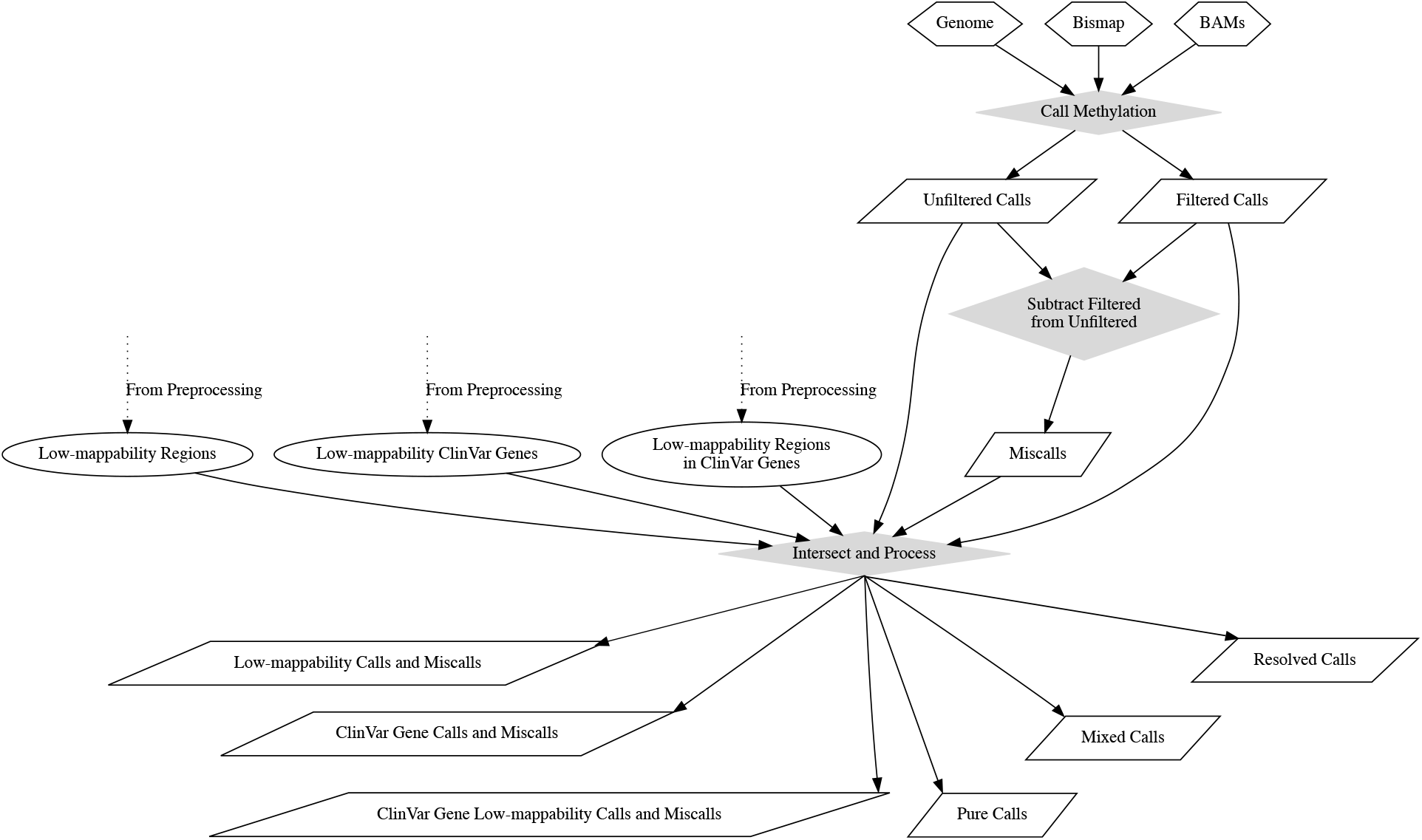
Detailed description of steps for calling methylation and intersecting the resulting calls with the low mappability regions determined in the preprocessing steps. Ovals are intermediate files. Gray rhombuses are processing steps with multiple arguments. Parallelograms are output files. Dotted lines indicate inputs that come from the preprocessing steps in Figure S.2. Parallelograms are output files.

The ClinVar VCF and GENCODE GFF were combined using bedtools and BEDOPS into a single BED file listing all GENCODE annotation regions that overlap one or more ClinVar variants (these regions will be referred to as “ClinVar regions”), which will be called the “ClinVar regions BED”. Each region in the ClinVar regions BED was expanded 2 kb upstream using bedtools slop to capture methylation upstream of the gene. The Bismap bedGraph (which did not contain zeroes) and an FAI index of the GRCh38 reference genome FASTA file were processed with bedtools, awk, and GNU sort to obtain a bedGraph containing all mappability data, including zeroes. This file will be referred to as the “Bismap complete bedGraph”. It was then combined with the ClinVar regions BED using bedtools map to create a file containing the minimum and mean mappability for each gene with a ClinVar variant.

The Bismap complete bedGraph was then filtered using awk to produce a file containing only low mappability regions (mappability < 0.01). The file with minimum and mean mappability for every ClinVar region was filtered likewise on minimum mappability. The two resulting files (one of low-mappability regions, one of ClinVar regions with low minimum mappability) were combined to produce a file of all low mappability regions that are in ClinVar regions.

Alignments of 2×101-bp paired-end EM-seq and whole genome bisulfite reads were processed using MethylDackel (with and without the custom patch) using a minimum MapQ cutoff of 10 and the default settings mentioned above and combined with the file of all low mappability regions in ClinVar regions to produce a list of all methylation calls in low mappability ClinVar regions (this will be referred to as the “low mappability ClinVar calls file”). A low-mappability methylation call, as used here, is defined as a methylation call that is in a region with Bismap mappability less than 0.01, as defined by the position of the C alone (positions of anchoring reads are not considered when determining this).

The low mappability ClinVar calls file was intersected with the file of ClinVar regions with low minimum mappability to produce a file of ClinVar regions with low mappability calls. The -wa option for bedtools intersect was used here, which writes a copy of the ClinVar region to the output file for each low mappability call in the region, in order that this file would contain multiple copies of each region, one per low mappability call in the region. These duplicates were then used to count low mappability calls by feeding the data to a custom Python script (found in the Nextflow script in the supplemental materials) which counted and combined the duplicates, producing a list of all ClinVar regions with low mappability calls and how many low mappability calls are in each region. A similar intersection was performed for all ClinVar regions with low minimum mappability (i.e. ClinVar regions that could potentially be affected) to produce a file of all calls (regardless of mappability of the C) in such regions, and a list of such regions with a count of calls per region.

Since this analysis was run for three different input masses and two sequencing protocols, the low mappability calls files were also processed through a custom Python script (see supplemental materials) to add a field specifying which input mass, sequencing protocol, and MethylDackel filtering setting were used. As the input mass would have been difficult to parse out of the BAM name due to inconsistent file name formatting, a CSV was created mapping BAM file names to input masses and given as input to this step. The same field was added to the lists of all ClinVar regions with low mappability calls and counts described previously. We also produced similarly-annotated files for all methylation calls (regardless of mappability), all methylation calls in GENCODE genes (with low minimum mappability) which contained ClinVar variants, all methylation calls in low-mappability regions that were also in GENCODE genes which contained ClinVar variants, counts of calls in in GENCODE genes which contained ClinVar variants regardless of mappability, methylation calls that are either 0% or 100%(regardless of mappability), and methylation calls which are neither 0%nor 100% (also regardless of mappability).

To assess miscalls (i.e. which Cs are filtered out by per-read filtering), as well as under/over-filtering with respect to per-C filtering (discussed above), and resolution of mixed methylation calls to 0% or 100%, pairs of these files were compared using bedtools to produce files showing the difference between filtered and unfiltered data. All files were examined and compared in Tableau^®^.

To compare the behavior of Bismark with bwa-meth, this analysis was re-run with the Bismark aligner using default settings and deduplicated using deduplicate_Bismark according to the documented protocol [18]. The name of the aligner was added to the field specifying the input mass, protocol, and MethylDackel filtering settings present in all output files containing methylation calls or counts of such calls in ClinVar regions (using the same custom Python script).

To compare the need for and the effect of mappability filtering on the more complete T2T v1.1 reference genome [26] (hereafter referred to as “T2T”), the analysis was re-run using the T2T reference in place of GRCh38 (supplemented with phage T4, phage lambda, phage Xp12, and pUC19 just as with GRCh38). To use the ClinVar and GENCODE datasets, which were respectively originally obtained as VCF and GFF files containing genomic coordinates in GRCh38, with the T2T reference, the positions were transferred over to the T2T reference using CrossMap [39]. This lift-over was done in two steps with separate chain files. First, the GRCh38 regions were lifted over using CrossMap to T2Tv1.0 using a chain file published on GitHub by Nico Alavi [1]. Then, the lifted-over T2Tv1.0 annotations were lifted over again to T2Tv1.1 using a chain file from the T2T Consortium [23]. Specifically, the ClinVar regions BED, with the ClinVar and GENCODE data already combined, was lifted over. This converted file was used in place of the ClinVar regions BED for analysis with the T2T reference. As there was no Bismap mappability file for the T2T reference, we downloaded the Bismap tool and generated a *k* = 100 multi-read Bismap mappability dataset for this reference genome.In total, this analysis was run on all combinations of sequencing method (WGBS or EM-seq), aligner (Bismark or bwa-meth), input mass (10, 50, or 200 ng), filtering (read-based filtering or no read-based filtering), and reference genome (GRCh38 or T2T), using a Nextflow[7] pipeline (see supplemental materials) which managed the execution of all the analysis tools needed.

### External Analysis Resources

Individual tools used in the analysis include sambamba[37], bigWigToBedGraph[17], MethylDackel, awk, python, GNU sort, GNU head, bedtools[32], and BEDOPS[25]. Versions of all tools used are specified in the nextflow 21.04.0 scripts using conda[24] dependency resolution.

The custom Python script that adds the input mass, sequencing protocol, MethylDackel filtering setting, and aligner name can be found at https://github.com/nebiolabs/low_bismap_methyl_calls.

The pull request for the patch adding mappability support to MethylDackel can be found at https://github.com/dpryan79/MethylDackel/pull/80. It was merged into MethylDackel in version 0.5.0.

The BBM format definition is available at https://github.com/dpryan79/MethylDackel/blob/master/BBM_Specification.md.

**Figure S.4:**
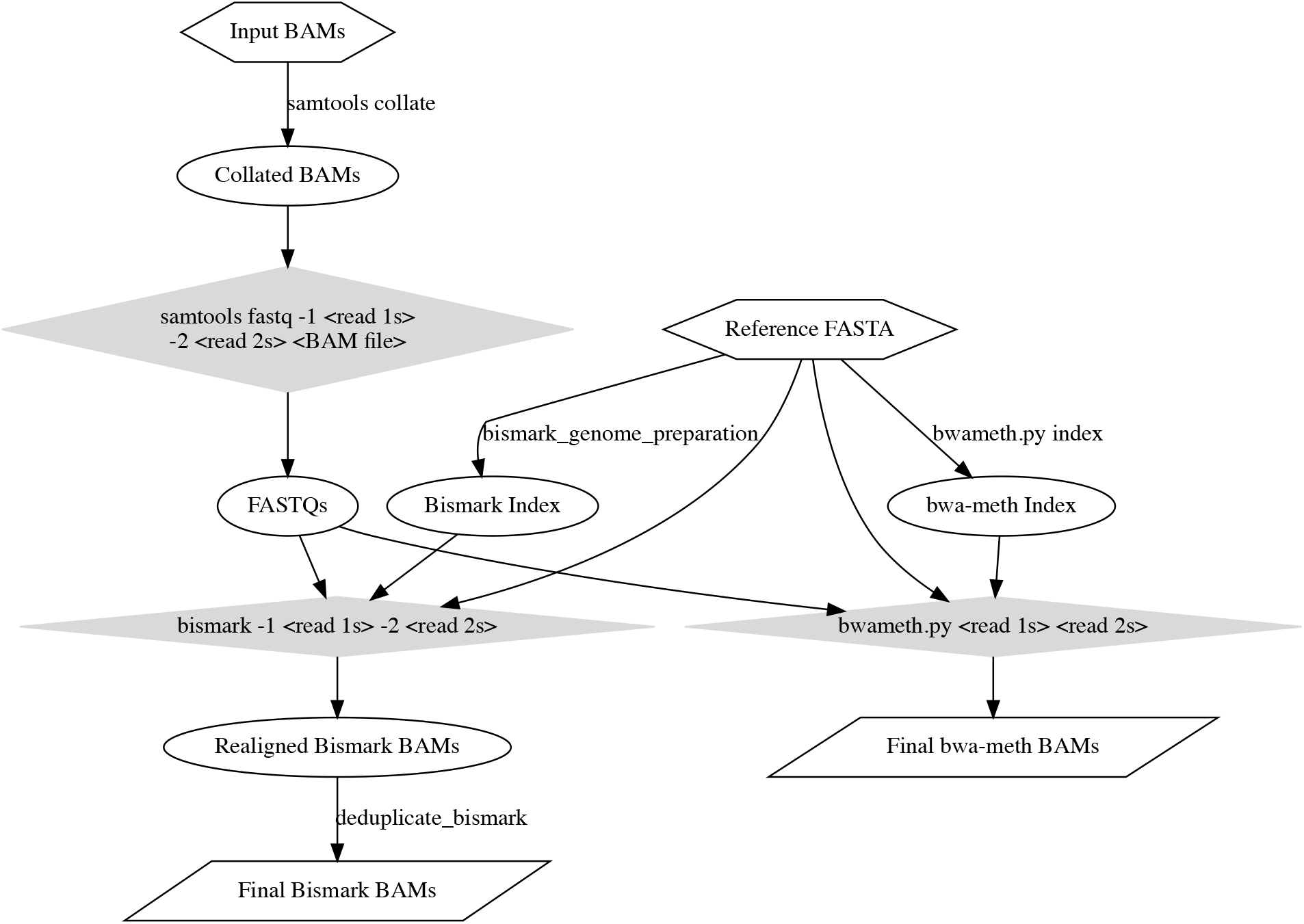
Detailed description of BAM realignment steps. Hexagons are input files. Ovals are intermediate files. Gray rhombuses are processing steps with multiple arguments. Parallelograms are output files. Labels on edges are processing steps with one input and one output.

The Bismap file used for this analysis was downloaded from https://www.pmgenomics.ca/hoffmanlab/proj/bismap/trackhub/hg38/k100.Bismap.MultiTrackMappability.bw

The ClinVar VCF was the July 22, 2019 version of ClinVar’s variants VCF, with a file name of Clin-Var_20190722.vcf.gz

The chromosome ideogram figure was generated using karyoploteR[10].

## Acknowledgments

We are grateful to the many people who contributed to the EM-seq project and made these libraries available. This work would not have been possible with out access to an extensible, open-source methylation caller. Many thanks to Chaithanya Ponnaluri for critical reading of this manuscript, to Matt Campbell and Ariel Erijman for helpful discussion, and to the organizers and supporters of the summer internship program at New England Biolabs that enabled this project.

